# Compositional shifts in the root microbiota track the life-cycle of field-grown rice plants

**DOI:** 10.1101/166025

**Authors:** Joseph Edwards, Christian Santos-Medellín, Zachary Liechty, Bao Nguyen, Eugene Lurie, Shane Eason, Gregory Phillips, Venkatesan Sundaresan

**Author notes:** **Corresponding author information:** Venkatesan Sundaresan, Plant Biology Department, UC Davis, Life Sciences Addition, 1 Shields Ave. Davis, CA 95616.

## Abstract

Bacterial communities associated with roots impact the health and nutrition of the host plant. While a multitude of static factors are known to influence the composition of the root-associated microbiota, the dynamics of these microbial assemblies over the plant life cycle are poorly understood. Here, we use dense temporal sampling of spatial compartments to characterize the root-associated microbiota of field grown rice (*Oryza sativa)* over the course of three consecutive growing seasons and two sites in diverse geographic regions. The root microbiota was found to be highly dynamic during the vegetative phase of plant growth, then stabilizes compositionally for the remainder of the life cycle. Bacterial taxa conserved between field sites can be used as predictive features of rice plant age by modeling using a random forests approach. The age-prediction models were used to reveal that drought stressed plants have developmentally delayed microbiota compared to unstressed plants. Further, by using genotypes with varying developmental rates, we show that shifts in the microbiome are correlated with rates of developmental transitions rather than age alone, such that different microbiota compositions reflect juvenile and adult life stages. These results suggest a model for successional dynamics of the root-associated microbiota over the plant life cycle.

## Introduction

Plants assemble soil derived root-associated microbial communities consisting of thousands of different species and strains (1–3). Taxa within the root-associated microbiota have been found to be beneficial for plant growth and resistance to biotic and abiotic stresses (4, 5). The root-associated microbiota can be partitioned into three spatially distinct root compartments with significantly different compositions (6): the soil adjacent to the root (the rhizosphere), the root surface (the rhizoplane), and the root interior (the endosphere) (6–8). It was found that each of these root-associated compartments have significantly different microbiota profiles from the communities in unplanted soil, thus indicating the roots enrich for subsets of the soil microbiota. This enrichment process and how the root-associated microbiota change throughout the lifecycle of the plant remain poorly characterized.

High resolution longitudinal sampling has allowed for characterization of the human gut microbiota successional progressions from infancy into adulthood (9–11). Newborns are characterized by a low diversity gut microbiota that increases with age. The composition of the human gut microbiota changes rapidly during the first three years of life, then stabilizes to become more like an adult microbiota after three years and these shifts correlate well with dietary intake. These patterns were consistent across three distinct human populations. Abnormal development of the human gut microbiota has been identified as a causal factor in the malnutrition disease kwashiorkor and treatment for the disease through diet amendments not only restores the weight of the malnourished children to levels comparable to healthy children, but also restores the development of the gut microbiota (12, 13). Furthermore, assembly of a normal gut microbiota has been implicated in recovery from infection by enteropathogenic bacteria (14). Thus, normal development of the gut microbiota is important for maintaining nutritional status as well as avoiding and recovering from enteropathogen infection.

By comparison, limited information is available about the spatiotemporal dynamics of the plant root-associated microbiota, especially for the root endosphere. Previously, we characterized temporal progressions of the microbiota across the rhizosphere-endosphere continuum of greenhouse grown rice seedlings from transplantation to two weeks after transplantation. Over this relatively short period of time the microbiota shifted to become more similar to those hosted by rice plants that were 42 days old, suggesting that plant age is an important factor contributing to the composition of the root microbiome. From this study, it was unclear how microbiota shift over larger intervals of time, especially across major developmental transitions. Dombrowski et. al found that both the rhizosphere and root microbiota of the perennial plant *Arabis alpina* were significantly different during 3 time points encompassing a 28-week period when grown in pots under greenhouse conditions (15). However, an early flowering mutant had no significant effect on microbiota compared to the non-flowering wild type plants, suggesting that soil residence time, rather than plant development stage, might be responsible for the observed shifts. Due to the relatively sparse time sampling employed in that study, it is not clear what patterns of change can be expected over the lifecycle of a plant when grown under field conditions, especially for annual plants which encompass most staple food crops. Specifically, it is unknown how root-associated microbiota vary on weekly timescale and whether the microbiota assemble in a consistent manner across geographically and climatically distinct regions. In this study, we use a high resolution spatiotemporal approach to detail the successional progression of the microbiota across the rhizosphere, rhizoplane, and endosphere compartments over the life cycle of rice plants grown under field conditions over multiple seasons and in two distinct geographically distinct regions of USA.

## Results

### Experimental Design and Sequencing

We first were interested in monitoring how root-associated microbiota shift throughout the life cycle of the rice plant under field conditions and whether changes in community profiles were consistent across multiple seasons and growing regions. To do this, we collected the rhizosphere, rhizoplane, and endosphere microbiome samples from rice plants grown in a submerged rice field in Arbuckle, California over the 2014 and 2015 seasons as well as the rhizosphere and endosphere samples of rice plants growing in a submerged rice field near Jonesboro, Arkansas in 2016. In the 2014 and 2015 seasons, the field was water seeded with *Oryza sativa* (cultivar M206, a commercially grown variety adapted to California). In 2014, the field was sampled weekly until harvest, a total of 19 weeks with each time point consisting of 8 replicates. In 2015, because we were interested in the initial root colonization stage, we sampled every other day for the first 4 weeks of growth, thereafter sampling every other week until harvest. Each time point in 2015 consisted of 4 replicates. For each season, we collected bulk soil (i.e. soil from the submerged field where no rice roots were present) until extensive root growth of the rice plants made it impossible to locate soil that was not proximal to any roots; this point was reached at about 8 weeks for the 2014 season and at 6 weeks for the 2015 season. For the 2016 season in Arkansas, we grew 2 early flowering rice genotypes: Sabine a tropical japonica variety cultivated widely across the Mississippi Delta region and CLXL745, a hybrid rice variety grown in Arkansas. We sequenced the V4 region of the 16S rRNA gene to determine the bacterial and archaeal component of the microbiome. After removing sequences classified as belonging to the host plant and removing OTUs that were not in at least 5% of the samples, we retrieved a total of 34,987,084 reads belonging to 10,893 OTUs across 1,111 samples.

### Microbiomes are shaped by distance from the root, growing regions, and plant age

To understand the underlying driving forces of microbial community variation in our data, we used Principal Coordinates Analysis (PCoA) of Bray-Curtis distances. We found that samples from 2014 and 2015 California samples as well as the 2016 Arkansas samples display a spatial pattern of divergence along the first principal coordinate, where communities in bulk soil samples cluster on one end of the axis and endosphere communities cluster on the other (Fig. 1A). PERMANOVA of pairwise distances between microbial communities indicated that the microbiota differed significantly between root-associated compartments (R2 = 0.22, P < 0.001), in agreement with our previous studies on the rice root microbiota. The microbiome samples from Arkansas and California separated along the second principal coordinate (Fig. 1B). This observation was supported by the PERMANOVA statistic (R2 = 0.13, P < 0.001).

**Figure 1.**
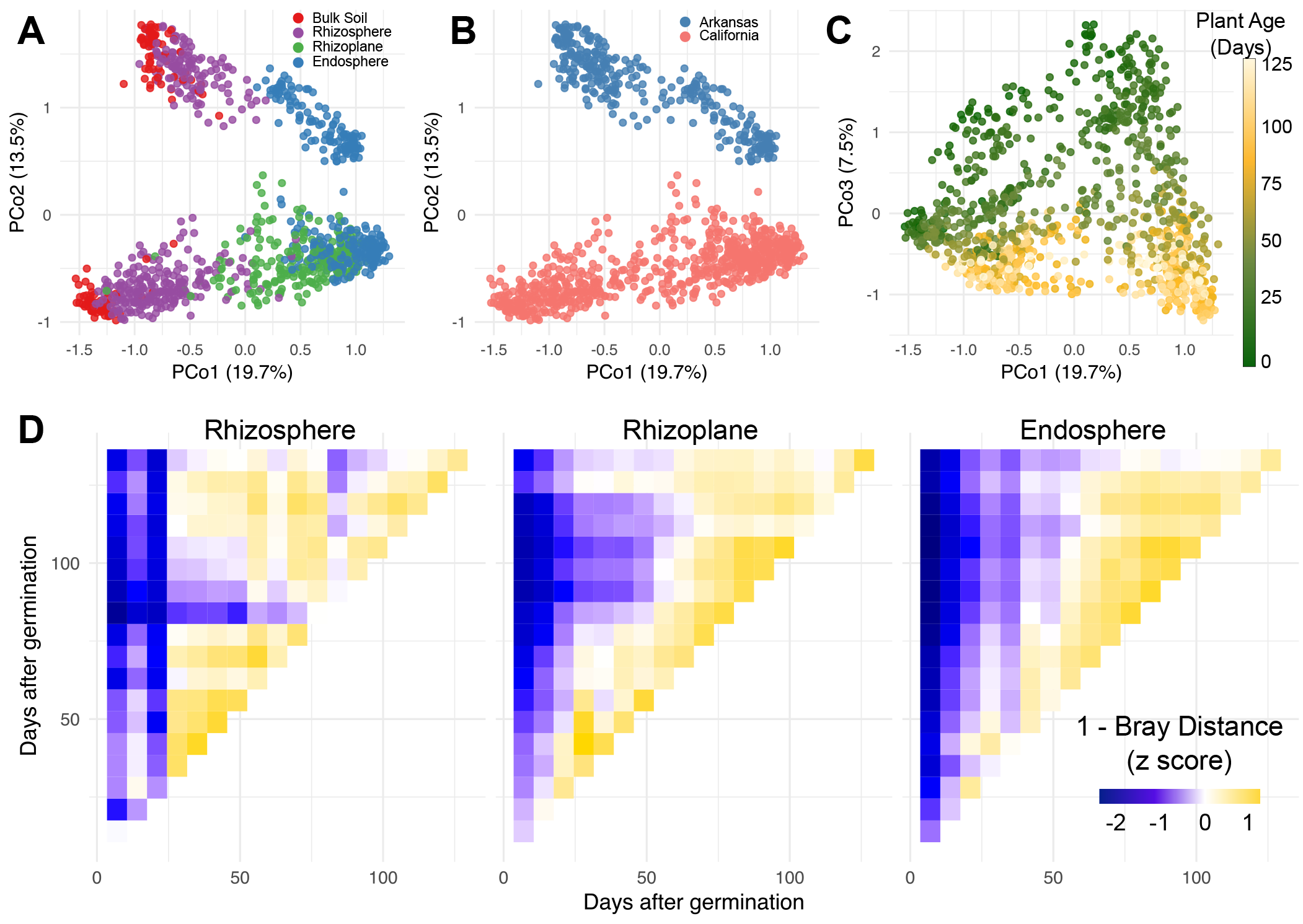
The root-associated microbiota stabilizes after 8-9 weeks after germination. **A)** Principal coordinate analysis (PCoA) of Bray-Curtis distances between samples colored by root-compartment. **B)** The same plot as in A but colored by the field location. **C)** The same analysis as A) and B), but now showing PCo1 vs PCo3 and the points are colored by the age of the plants from which the samples were taken **D)** Heatmaps showing mean pairwise z-scores for Bray-Curtis similarity between time points in each compartment for the 2014 California samples.

We next measured the effect of plant age on the root-associated microbiota. We found that this effect is observable across the third principal coordinate of the PCoA plot and is consistent across the two growing regions, and across the two growing seasons within California (Fig. 1C). The amount of variance partitioned to the effect of plant age on the root-associated microbiota was comparable between the two growing regions (California: R2 = 0.093, P<0.001; Arkansas: R2 = 0.080, P<0.001). We observed that the rhizoplane and endosphere microbial communities shift over the first seven to eight weeks after germination, but stabilize thereafter (Fig. 1D, Sup. Fig. 1). This pattern was observed over both growing seasons in California and also in the Arkansas field trial. However, a stabilization pattern was not observed in the rhizosphere microbial communities. It is interesting to note that the selected rice varieties grown in California and Arkansas reach panicle initiation (entry into reproductive growth) 8-9 weeks after germination in their respective locations, suggesting a correlation between plant developmental stage and root-associated microbiota dynamics.

The multi-year sampling scheme of our experimental design allowed us to quantify the effect of year to year variation on the root-associated microbiota. Although the root-associated microbiota varied significantly across the two different years, the effect was small (R2 = 0.007, P = 0.001) compared to the other factors analyzed within this experiment. Together, these data suggest that the root-associated microbiota shifts in each root-associated compartment during the vegetative growth stage of the season and stabilizes upon entry into reproduction and that these patterns are consistent across different growing seasons.

### Microbiota dynamics over the season are marked by increasing and decreasing shifts in relative abundance of specific phyla

We next sought to characterize the specific phyla responsible for the significant differences between the root-associated compartments and how these various phyla change in abundance over the course of the season. Overall, we noticed similarities in patterns in phyla abundance over time between the two growing regions (Fig. 2A). Because the Proteobacteria phylum contains a broad phylogenetic makeup and because Proteobacteria make up a large proportion of the rice root-associated microbiota we further divided the Proteobacteria phylum into its respective classes for this analysis. To model increasing or decreasing relative abundance of individual phyla between the various root-associated compartments, we assigned each compartment a value relative to its spatial position: bulk soil was position 0, rhizosphere was position 1, rhizoplane was position 2, and endosphere was position 3. We modelled how each phylum either increased or decreased over these positions using beta regression. Beta regression is used for modeling dependent variables which lie in the interval (0, 1) and is thus useful for modelling individual taxa as a relative proportion of the total microbial community.

**Figure 2.**
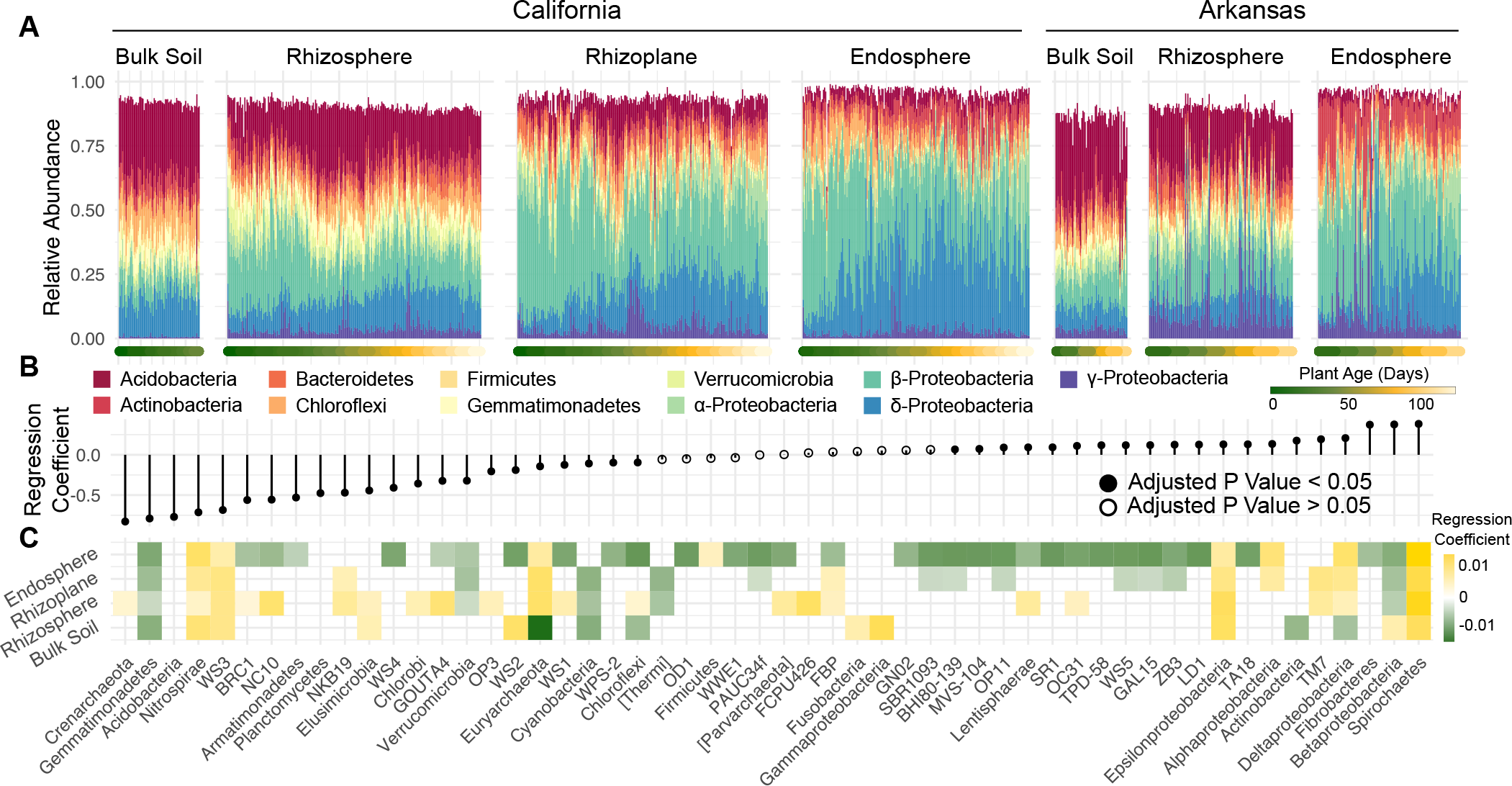
Shifts in the microbiota over time are associated with increasing and decreasing phyla. **A)** Bar plots of the top 11 phyla abundance over the course of the seasons in each compartment. Each bar represents 1 sample that was taken throughout the course of the growing season. The bars are ordered by the age of the plant as indicated by the colored points beneath each bar. Both the 2014 and 2015 data were used for this graph. **B)** Beta regression coefficient estimates for microbial phyla that are either increasing (above 0) or decreasing (below 0) in relative abundance from the outside of the root to the inside of the root. **C)** Beta regression coefficient estimates for microbial phyla that are increasing (above 0) or decreasing (below 0) in relative abundance over the course of the seasons in each compartment.

Using this method, of the 55 detectable phyla, we identified 42 phyla which significantly differed in spatial distribution from the exterior to the interior of the root (Fig. 2B). There were 20 phyla whose relative abundance significantly increased from the soil environment to the root interior, and 22 microbial phyla whose relative abundance decreased from the soil environment to the root interior. The absolute values of the regression coefficients were overall greater for root-depleted phyla, suggesting that a smaller proportion of the soil microbiota is enriched by the plant while more microbial taxa are depleted.

We again performed beta-regression within each compartment to identify phyla that significantly increased or decreased in relative abundance as a function of plant age (Fig. 2C). In general, trends in abundance over the season were consistent across the rhizocompartments (excluding the bulk soil) for each phylum. For instance, in the rhizosphere, rhizoplane, and endosphere Betaproteobacteria, Verrucomicrobia, and Gemmatimonadetes consistently decreased over the course of the season, while Deltaproteobacteria, Epsilonproteobacteria, Euryarchaeota, and Spirochaetes all increased. In the rhizoplane and endosphere, most of the taxa with dynamic abundance patterns significantly decreased over the course of the season (12/22 and 32/40, respectively), suggesting that the root-associated compartments are initially colonized by a diverse set of microbes from the soil, but are either eventually outcompeted by other taxa or are selected against by the host plant. Together, these results indicate that despite large differences in microbiota composition between the growing regions, consistent trends in abundance at the phylum level define microbiota differences between the rhizocompartments and over the course of the growing season.

### Plant age is predicted by subsets of the microbiota that are conserved across field sites

We next sought to identify individual microbial OTUs which could be used to discriminate plant age. To do this, we began by developing compartment-specific full Random Forests (RF) models for both the endosphere and rhizosphere samples by regressing the relative abundance of all OTUs against the age of the plants from which the samples were collected. For our training data we selected samples from the California 2014 and Arkansas 2016 data. Half of the samples from each time point and rhizosphere and endosphere compartments were used for the training set. From these full models, we were able to rank individual OTUs by their importance in contributing to the accuracy of age prediction by the models. Because not every OTU included in the full RF models will contribute the models’ accuracy, we next performed 10-fold cross validation while simultaneously removing less important OTUs to evaluate model performance as a function of inclusion of the top age-discriminant OTUs. We found that there was minimal increase in accuracy when including more than 85 of the most important OTUs (Supplemental Fig). We next developed sparse RF models for the rhizosphere and endosphere compartments using the 85 most important OTUs from each full RF model.

We found the sparse RF models explained 91.5% and 88.4% of the variance related to plant age for the rhizosphere and endosphere, respectively. Plant age was accurately predicted by the compartment-specific sparse RF models for the California and Arkansas sites (Fig. 3A). Additionally, we note that the ages of plants sampled during from California during the 2015 season were able to be accurately predicted, despite the models not being trained on this data. These results indicate that specific sets of microbes in both the endosphere and rhizosphere behave in consistent patterns over the course of the rice plant’s life cycle between seasons and across geographic regions.

**Figure 3.**
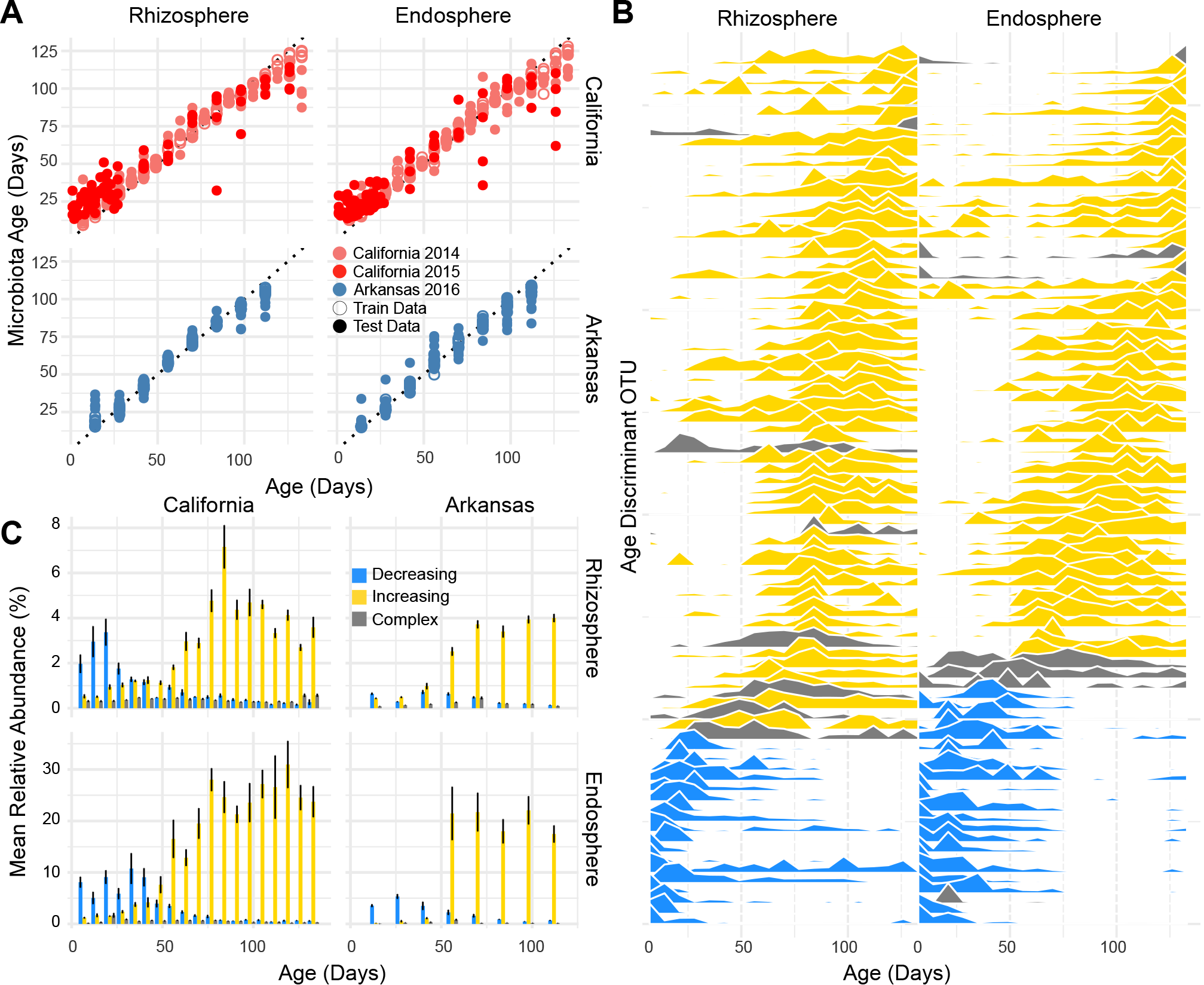
Random forests model detect taxa that are accurately predictive of plant age. **A)** The result of predicting plant age using the sparse RF models for the 2014 and 2015 season. Each point represents a predicted age value. **B)** Abundance profiles for the age discriminant OTUs in rhizosphere and endosphere compartments over the course of the California 2014 growing season. OTUs are ordered along the y-axis by timing of peak abundance. The orders of the OTUs on the y-axis are not shared between rhizosphere and endosphere, despite both the models for each compartments sharing a subset of age-discriminant taxa. OTUs are colored by their classification as increasing, decreasing, or complex over time (see color scale in panel C). See table for order of OTUs. **C)** Mean total abundance for the age-discriminant taxa across sites and compartments.

We next examined the phylogenetic composition of the 85 OTUs used for plant age prediction in the rhizosphere and endosphere sparse models. The OTUs used in the sparse RF models were phylogenetically diverse. In the endosphere model, the model used OTUs belonging to 10 phyla, 19 classes, 29 orders, and 37 families, while the rhizosphere model used OTUs belonging to 11 phyla, 24 classes, 37 orders, and 45 families. Consistent with general trends in alpha diversity between the compartments, the rhizosphere model contained a more phylogenetically diverse set of microbes for classification. We used a linear model approach to classify the OTUs included in each model as having increasing, decreasing, or complex abundance patterns over the course of the season (Fig. 3B). We found that for both the rhizosphere and endosphere sparse RF models most of the OTUs increased in abundance over the season, while fewer OTUs were classified as complex or decreasing in abundance over the season (Fig. 3B). The rhizosphere and endosphere models shared 22 OTUs, 20 of which were in agreement for increase/decrease/complex classification. Within the OTUs classified as increasing or decreasing for the rhizosphere and endosphere models, Betaproteobacterial OTUs were the most represented class (Fig. 3B). At the order level, however, the OTUs making up the increasing or decreasing fraction within the sparse models were significantly different (Sup. Fig. 2). Betaproteobacterial OTUs decreasing in abundance were mainly composed of Burkholderiales, while the increasing OTUs were mainly composed of Rhodocyclales and SBla14. We also noticed differences between the rhizosphere and endosphere RF models for the taxa they were using for classification. The endosphere model used many more Alphaproteobacterial OTUs than the rhizosphere model, which mirrors differences in trends at the phylum level (Fig 2). Similarly, the rhizosphere model used OTUs from the phylum Verrucomicrobia, whose relative abundance at the phylum level is significantly higher in the bulk soil and rhizosphere than the endosphere (Fig. 2B).

We next inspected how much of the total microbiota the rhizosphere and endosphere age-discriminant taxa compose over the course of the season (Fig. 3C). Overall, the age-discriminant OTUs composed a greater proportion of reads in the endosphere compared to the rhizosphere. This was consistent across both the California and Arkansas sites. As expected, the OTUs that were decreasing in relative abundance as a function of rice plant age were dominant at the beginning of the season, while those identified as increasing in relative abundance were dominant at the end of the season. Interestingly, we noticed that the switch in dominance between increasing/decreasing classified OTUs occurred 8 to 9 weeks after germination, coinciding with the switch from vegetative to reproductive growth for the included varieties.

We next asked if the early colonizing age discriminant OTUs were significantly enriched in their respective root compartments compared to unplanted soil controls. We were only able to use the early time points (less than or equal to 49 days after germination) as a comparison due to the inability to gather bulk soil controls in the latter part of the season. We found that the microbes in the RF models classified as decreasing in abundance were predominantly enriched in the rhizosphere and endosphere compared to bulk soil controls (Sup. Fig. 3). The microbes classified as increasing had more similar relative abundance patterns to the bulk soil control samples. Because this analysis was constrained to using the earlier time points, we expect that the increasing OTUs would be predominantly enriched compared to the bulk soil controls in the latter time points. These results suggest that the early colonizing OTUs included in both the rhizosphere and endosphere sparse RF models are selected either actively or passively by the plant roots and their abundance profiles are likely not a product of edaphic processes separate from the plant.

### Drought stress is associated with delayed development of the endosphere microbiota

Drought is one of the most common and devastating stresses to affect rice production around the world. While soil microbes have been implicated in alleviating drought symptoms in various plant species under laboratory conditions (16, 17), it was unknown until recently how water deprivation affects the rice root-associated microbiota. Drought stress most strongly alters the endosphere microbiota (relative to the rhizosphere and bulk soil) with Actinobacteria and Chloroflexi strongly increasing in abundance and Deltaproteobacteria and Acidobacteria strongly decreasing under drought conditions. Despite large changes in the rice microbiota in the face of drought stress, it is unknown how these changes reflect on normal development of the microbiota.

To study how drought stress may affect normal development of the microbiota, we used our sparse age-predicting RF models in conjunction with the data collected from Santos-Medellín et al. to model plant age as a function of the microbiota (Fig. 4A). Santos-Medellín et al. analyzed the rhizosphere and endosphere microbiota of 49-day old drought stressed and well-watered rice plants growing in 3 diverse soils under greenhouse settings (18). When predicting ages from the samples included in this experiment, we found that the soil in which the plants were growing had the largest effect on variation in age predictions (F = 55.69, P < 2x10^-16^ ANOVA. The variation due to soil source was largely caused by samples originating from the Davis soil being predicted as younger compared to samples from plants grown in Arbuckle and Biggs soil. This result is not surprising because the Arbuckle and Biggs soil have cultivated rice every summer season for the previous 8 years while the Davis field has been fallowed for the previous 6 years. Thus, it is likely that soil cultivation history can affect the accuracy of the age-predicting RF models.

**Figure 4.**
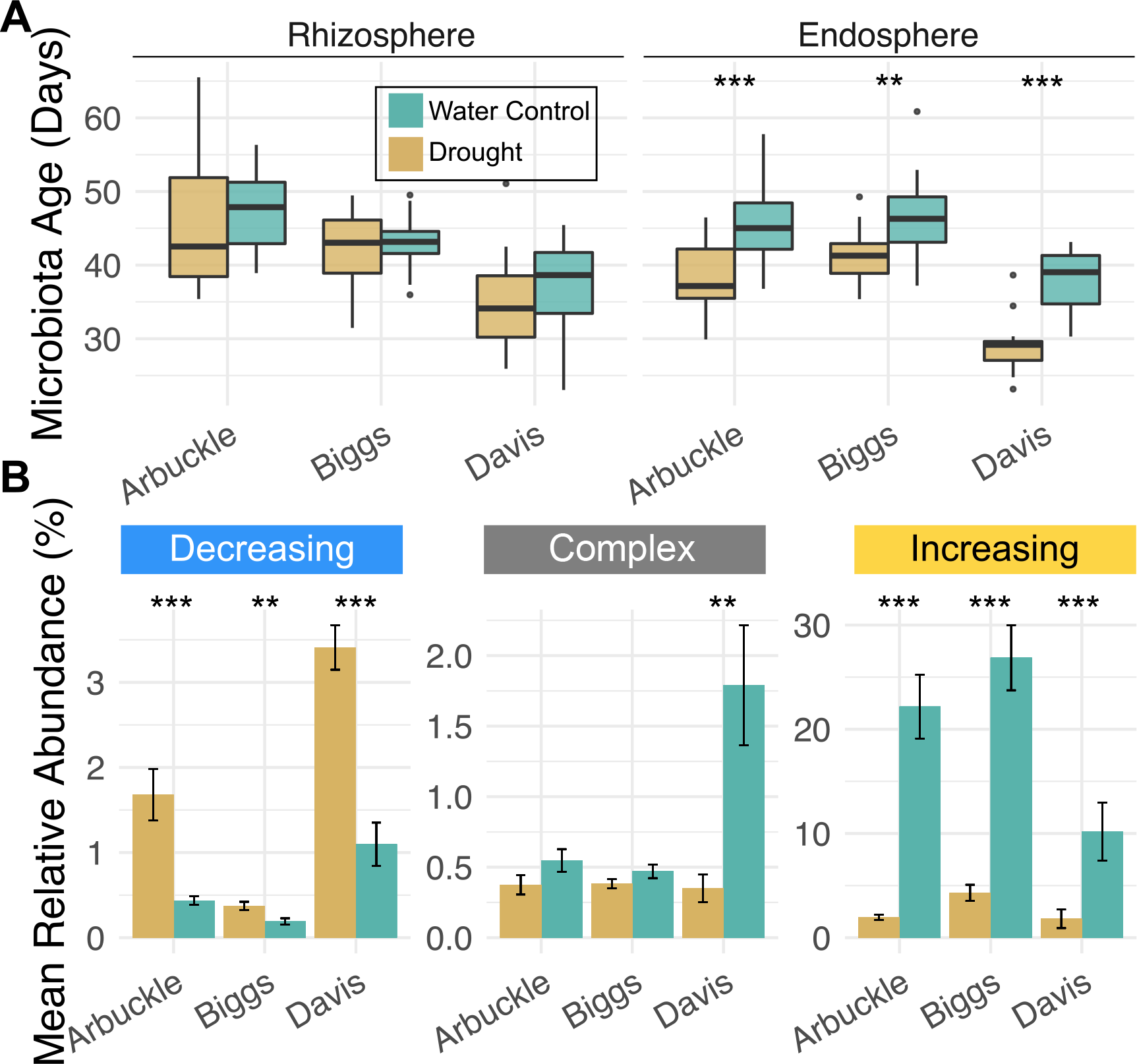
Drought exposure is associated with delayed development of the endosphere microbiota. **A)** Microbiota age predictions for rhizosphere and endosphere samples from well-watered and drought exposed plants. **B)** Abundance of age discriminant OTUs in the endosphere samples of well-watered and drought exposed plants. Color scales are shared between A and B.*** P < 0.001, ** P < 0.01, * P <0.05.

Watering treatment had the second largest effect on plant age prediction (F = 32.54, P = 4.72x10^-8^ ANOVA). This effect was only apparent in the endosphere (adjusted P = 7.9x10^-9^, Tukey’s post hoc test) and not in the rhizosphere (adjusted P = 0.34, Tukey’s post hoc test). This finding is consistent with the conclusions by Santos-Medellín et al., that the endosphere microbiota is most effected by water deprivation compared to the other rhizocompartments. In each soil, the drought stressed plants hosted endosphere microbiota that were consistently less mature than well-watered plants. We asked whether this was an effect of the early-colonizing microbes reemerging and becoming more abundant in the endosphere of plants exposed to drought or whether the late emerging microbiota was declining in abundance compared to well-watered plants (Fig. 6B). We found that both instances occurred: the early colonizing microbiota were significantly more abundant and the late emerging microbiota was significantly reduced in drought exposed endospheres compared to well-watered controls. Together these data indicate that drought stress is associated with delayed development of microbiota in the root endosphere, but not rhizosphere.

### The root-associated microbiota of distant field sites converge in similarity during the growing season

Despite clear distinctions in microbiota composition between the growing regions, PERMANOVA indicated a significant interaction between plant age and the difference in microbiota composition between the sites (R2 = 0.018, P < 0.001). We found that both the endosphere and rhizosphere microbiota became significantly more similar over time between the two field sites (Fig. 5A). Compared to the rhizosphere, the endosphere microbiota started the season as more dissimilar between the sites, but reached comparable levels of similarity by the end of the season. This trend did not hold for the bulk soils (Sup. Fig. 4): the bulk soil communities between the sites did not show any change in similarity over time (P =0.1), although we note that we were only able to sample bulk soil microbiomes over a 5-week span at the beginning of the season due to root proliferation later in the season. Interestingly, the similarity in microbial composition between the plants growing in each site stabilized after the rice plants had entered the reproductive phase. This result suggests that plants growing in disparate field sites initially acquire divergent root-associated microbiota, but the composition of these communities begins to converge throughout vegetative growth and maximizes and stabilizes during the reproductive phase. Together these results suggest a host selection within the rhizosphere and endosphere that acts on similar microbes within the microbiota provided by the soil community.

**Figure 5.**
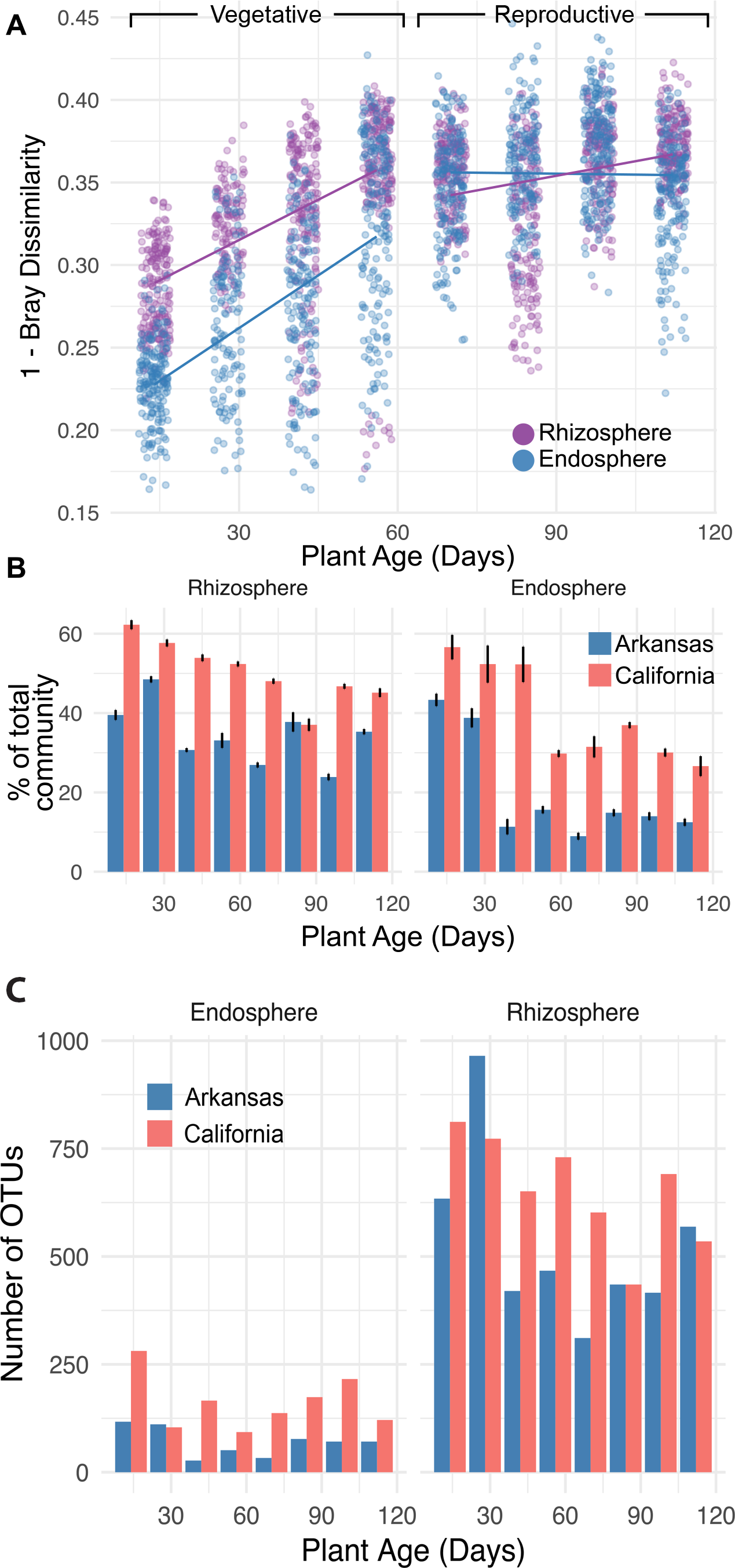
Rhizocompartments become more similar between field sites as a function of plant age. **A)** Pairwise distances between each site within each common time point and each common compartment. **B)** Number of OTUs defined as site-specific in each time point. Colors indicate which soil the OTU was found to be enriched in. **C)** Mean total relative abundance of the site specific OTUs within each time point.

Additionally, _the composition of the plant age-predicting RF compartments was largely composed of OTUs whose abundance were increasing over the season at each field site. We hypothesized that this was due to the composition of the earlier time points being largely composed of site-specific OTUs. To test this hypothesis, we first identified OTUs in each time point whose abundance was skewed towards a specific site using edgeR, a software package that incorporates empirical Bayes methods to account for overdispersion common across count data (19). After correcting for multiple comparisons, we identified 3218 OTUs in the rhizosphere and 865 OTUs in the endosphere that were significantly skewed towards one of the field sites. We did not notice a clear trend in the number of OTUs called per time point (Sup. Fig. 5). When taking the overall relative abundance of the site-specific OTUs into account, however, we found that the site-specific OTUs made up a significantly greater proportion of the total microbiota of the earlier time points than the later time points in both field sites and both the endosphere and rhizosphere compartment (Fig. 5B, table). This effect was different across the rhizosphere and endosphere, with the endosphere having a more pronounced effect. We suggest that this difference might be due to the rhizosphere hosting plant-responsive microbes as well as soil microbiota which do not respond to plant stimuli. These results suggest that the initial colonization of the rhizosphere and endosphere is largely composed of site-specific OTUs; however, later in the season, a set of OTUs conserved between the two field sites begin to establish in the endosphere and rhizosphere, but not in the bulk soil.

### Developmental stage is a driver of microbiota succession

Our sparse RF models appeared to detect progressions in the root-associated microbiota that correlate with developmental transitions in the rice plants. However, plant developmental transitions also co-vary with climatic and edaphic factors which may have indirect effects on the microbiota, making these factors difficult to uncouple from the direct effect of plant developmental stage on the root-associated microbiota. Because our California study was confined to one variety and our Arkansas study was confined to studying two varieties which have very similar developmental progressions, we were unable to unambiguously discriminate the effect of plant development from that of environmental factors on root-associated microbiota assembly. To investigate the direct effect of plant developmental stage on the root-associated microbiota, it is necessary to be able to distinguish the effect of plant age from developmental progression. With this aim, we grew 4 rice varieties of the same temperate *japonica* group of *Oryza sativa* at the same California field site during the 2016 season: Kitaake, California varieties M206 and M401, and Nipponbare. These cultivars grow with different developmental progression rates, (Fig. 6A). These varieties were water seeded in the California field in a complete randomized block design. We sampled plants within each plot every two weeks throughout the season, collecting rhizosphere, rhizoplane, and endosphere fractions from the plant roots. We also scored the plants for their developmental stages Our revised field design allowed us to also collect bulk soil samples throughout the entirety of the season as compared to the 2014 and 2015 seasons which restricted our soil sampling due to the invasiveness of the rice roots. After sequencing the V4 region of the 16S rRNA gene and filtering out plastid OTUs, we obtained 11,986,615 total sequences composing 10,547 OTUs from 469 samples.

**Figure 6.**
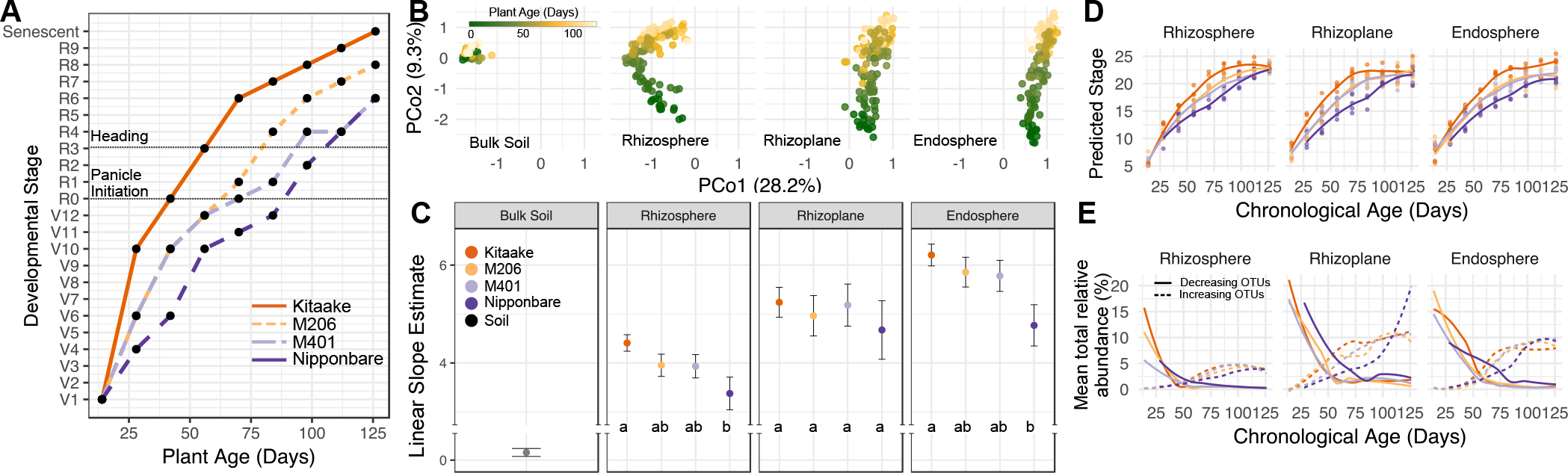
Varieties with different developmental rates have skewed microbiota progressions. **A)** The developmental stage of the tested varieties as a function of plant age. We staged each variety using descriptors previously described (53). R0 corresponds to the panicle initiation stage. It is important to note that M401 and M206 had nearly identical times to panicle initiation, but afterwards diverged in time to heading. **B)** PCoA of the 2016 data indicating root-associated compartment and plant age are major determinants of microbiota structure. **C)** Linear slope estimates for the principal coordinate 2 in panel B as a function of plant age for each variety and each compartment. A higher slope indicates that the microbiota is progressing faster along PCo2 as a function of plant age. **D)** The predicted developmental stage of the 2016 data as predicted by the stage discriminant sparse RF models. **E)** Total relative abundance estimates for increasing and decreasing stage-discriminant taxa between each cultivar and compartment.

The data from the 2016 season had similar trends as exhibited in the 2014 and 2015 seasons in California as well as the field trial in Arkansas. The root-associated compartments hosted distinct microbiota (Fig. 6B, R2 = 0.328, P < 0.001) and these microbiota varied significantly due to plant age (Fig. 6B, R2 = 0.068, P < 0.001). We found that genotype had a very small overall effect on the root-associated microbiota (R2 = 0.010, P < 0.001). This level of variance is smaller than what we have previously detected in rice, but is not surprising given that the included varieties were constrained to the temperate *japonica* group. We noticed that there was a significant statistical interaction between plant age and genotype (R2 = 0.008, P < 0.001), suggesting that the trends in microbiota shifts over the season differ depending on the genotype. It is important to note that we did not find a significant interaction in the Arkansas field trial between rice genotype and age, presumably because the included varieties were very similar in their developmental progression rates. Similarly, we found that plant developmental stage explained more variance in our dataset than plant age (R2 = 0.082, P < 0.001), again suggesting that developmental stage is an important descriptor for root-associated microbiota assembly. To further inspect this observation, we asked whether there was a significant interaction between plant genotype and plant age along the second principal coordinate (PC) of Fig 5B as a function of plant age. We used the second PC because it was the axis best differentiating plant age. We hypothesized if plant developmental rate were to have an effect on the root associated microbiota, then the faster developing varieties would have steeper slopes than the slower developing varieties. We found a significant interaction between plant age and genotypes across the second principal coordinate axis for both the rhizosphere and endosphere, but not the rhizoplane (P = 0.009, 0.007, 0.26, respectively), indicating that microbiota of the chosen cultivars mature at different rates. When inspecting the slope estimates for each variety’s microbiota over the second PC as a function of plant age, we observed that progression of the microbiota followed similar trends as the plant development rate variation between the cultivars (Fig. 6C). Specifically, Kitaake and Nipponbare had significantly different slope estimates in the rhizosphere and endosphere, while M401 and M206 had intermediate slope estimates that were not significantly different to the other varieties. Each compartment had a positive slope over the course of the growing season, but unplanted soil had a much more gradual positive slope than the rhizocompartments. Taken together these results indicate that root microbiota shifts throughout the season appear to arise independently of shifts in the bulk soil, and the rate of these shifts in the rhizosphere and endosphere correlate with the developmental progression rate of the host plant.

We next used the sparse RF models generated from the California and Arkansas data to predict the age of samples taken from the 2016 season. In general, the models predicted the ages of the plants accurately (Sup. Fig. 6A). The endosphere predictions showed significant variation due to cultivar (P = 0.001, ANOVA) while the rhizosphere predictions did not (P = 0.191). Furthermore, the endosphere model showed trends in the predictions that correlated with developmental progression rate variation between the cultivars with Nipponbare having predicted ages significantly younger than Kitaake and M206 (adjusted P = 0.0012 and 0.0021, respectively). When observing the total abundance of age discriminant OTUs from the models between the cultivars over time, we found that OTUs with increasing abundances in the endosphere took a significantly longer period of time to establish in Nipponbare compared to Kitaake and M206 (adjusted P = 0.0002 and P = 0.0016, respectively; Sup. Fig. 6B). These data suggest that the rhizosphere age-predicting model is largely robust to differences in phenology due to plant genotype, but that the accuracy of the endosphere model is affected by variation in plant developmental rate.

To have a better understanding of which microbes are associated with the various developmental stages, we formed new RF models for predicting plant developmental stage rather than plant age. Development in plants is a continuous process marked by numerous stage-specific morphological features. We monitored the developmental stage of the genotypes throughout the season assigning a numerical value from 1-27, with a value of 1 corresponding to a recently germinated seeding and 27 corresponding to a senescent plant. Panicle initiation corresponded to a value of 18, thus a plant that has entered the reproductive phase has a value of 18 or higher and a plant in the vegetative phase has a value of 17 or lower. We followed the same approach as previously mentioned to develop the RF models. Briefly, we trained full RF models in each compartment where we regressed the full dataset of OTUs against the developmental stage number for a training set of samples. From these full models, we sequentially removed OTUs of lower importance while performing 10-fold cross validation. We found that the models were near peak accuracy when including 54 of the most important OTUs for each compartment. With these 54 most-important OTUs, we developed sparse RF models that model the microbiota as a function of plant developmental stage (Fig. 6D). When the predicted values of plant developmental stage were plotted as a function of the plant’s chronological age, we found that the predictions accurately matched the developmental progressions that we witnessed in the field (see Fig. 6A). Variation in developmental stage prediction slopes were significantly affected by plant genotype in each compartment (Rhizosphere P = 8.1x10^-8^, Rhizoplane P = 3.3x10^-5^, Endosphere P = 2.3x10^-2^). These results indicate that OTUs within the microbiota in each compartment can be used to predict plant developmental stage, even when developmental progression rates differ between rice cultivars.

We again classified the OTUs included in each model to detect whether the OTUs had significantly increasing, significantly decreasing, or complex distributions throughout the season. The phylogenetic classification of increasing/decreasing OTUs mimicked those of the age-discriminant models (Sup. Fig. 2 and Sup. Fig. 7). Of the OTUs classified as decreasing over the season, 3 were shared between the compartment specific models, while 9 of the increasing OTUs were shared. We next sought to address whether OTUs included in the models peak in abundance at different times in the season for the different cultivars. To do this, we calculated the average abundance of each OTU included in each model at each time point for each cultivar. For every compartment, we found that the decreasing set of OTUs peaked later in Nipponbare than the other cultivars. When taking the total abundance of increasing/decreasing OTUs into account, we found that the decreasing OTUs tended to persist for a longer period of time in Nipponbare than the other varieties (Fig. 6E). Similarly, the OTUs classified as significantly increasing over the season took longer to establish in Nipponbare than the other varieties. This pattern was consistent across each compartment. We note that the developmental stage sparse RF models used fewer OTUs than the plant age models to make accurate predictions. Together, these results indicate that both plant age and developmental stage are important drivers of the root microbiota.

## Discussion

Microbial communities associated with plants can have strong positive and negative effects on plant health and nutrition (4) as well as contribute to greenhouse gas emissions (20, 21) and biogeochemical cycling (22). Understanding and studying the spatial distribution as well as temporal progressions of the root microbiota is therefore an important aspect for crop and environmental improvement. Our analysis provides a detailed characterization of the spatiotemporal dynamics of the root microbiota and reveals insights into how plant development and abiotic stresses affect development on the microbiota.

### The dynamic patterns of root-associated microbiota are consistent across field sites and seasons

Previous studies conducted with potted plants under greenhouse conditions have indicated that rhizosphere microbiota significantly varied across time points (23, 24) and one study has suggested that the endosphere microbiota is different across three time points (25). While these studies provided important evidence that the composition of plant microbiota is dynamic during plant growth, due to the limited extent of both temporal and spatial sampling it has not been possible to formulate a comprehensive picture of the changes in the microbiome during the plant life-cycle, as has been accomplished for humans. Additionally, the difference between greenhouse and field conditions is important given that nutrient dynamics and plant physiology vary under the two settings (26, 27) and it has been demonstrated that plant microbiota are also altered by field conditions (6). Similarly, importance of ecological factors driving community assembly are overestimated if the experimental observations are constrained to one soil type or season (28).

With these considerations in mind, we found that the microbiota in the rhizosphere as well as the rhizoplane and endosphere change in composition significantly over the life cycle of the host plant (Fig. 1). The changes in microbiota structure were consistent across multiple years of cultivation in the same field (Fig. 1A-C). Moreover, although we found large differences at the OTU level between the two tested field sites in California and Arkansas, at the phylum level we found that there were remarkable similarities in spatiotemporal profiles of microbiota abundance between the field sites despite large geographic distances, climatic differences, and cultivation practices (Fig. 2A). In general, there were more phyla that were decreasing in relative abundance in the endosphere over the life cycle of the rice plants, while fewer phyla were increasing in relative abundance. In the rhizosphere, we found that more phyla were increasing rather than decreasing over the life cycle (Fig. 2C). These data reinforce the exclusionary role of the endosphere compartment compared to the rhizosphere compartment as observed in other studies (6–8), but indicates that exclusion of microbes in the endosphere is age sensitive. Beta and Delta-Proteobacteria were the dominant classes enriched in the root endosphere compared to the bulk soil. These two phyla showed opposing patterns of abundance over the season: Betaproteobacteria was largely decreasing in abundance while Deltaproteobacteria increased (Fig. 2C). The human gut microbiome is also characterized by contrasting patterns of certain microbes: Enterobactericeae and Bifidobacteriaceae are early colonizers of the infant gut, while Ruminococcaceae and Prevotella become more prevalent later in development (11).

### The microbiota stabilizes 8-9 weeks after germination

Our high-resolution sampling scheme allowed us to deeply characterize the developmental patterns of microbiota over the course of the season. The rhizoplane and endosphere microbiota progressed over the course of the first 7 to 8 weeks after germination, but then stabilized in composition thereafter (Fig. 1D). This was consistent across the two years of sampling from the California field as well as for samples collected from the Arkansas field. These results resemble observations of the human gut microbiota dynamics. Clear compositional differences exist between the gut microbiota of human subjects from the US and subjects from Malawian and Amerindian villages (10). However, within each population, progression from an infant to adult microbiota configuration occurred around 3 years after birth. A shotgun metagenomic approach revealed that the adult and baby microbiota are functionally distinct, with age associated changes in vitamin synthesis and metabolism genes. The functional attributes of the stabilizing microbiota associated with plant roots remains uncharacterized. We find the similarity in stabilization patters surprising given that the mode of microbiota acquisition between mammals and plants is different (3), where plants mainly assemble microbiota by horizontal transmission from the surrounding soil microbiota and the mammal gut microbiota is acquired through a combination of vertical transmission from the mother during birth and breastfeeding and horizontal transmission during feeding or cohabitation.

### Creating a baseline for normal rice root microbiota progression

By employing a machine learning approach, we were able to model rice plant age as a function of fluctuating relative abundances of OTUs (Fig. 3). Using the random forests algorithm, we identified OTUs in the rhizosphere and endosphere compartments that were discriminant of plant age. Using these sets of OTUs we were able to accurately predict plant age of samples gathered from the California and Arkansas field sites. We were also able to use these models to accurately predict the age of rice plants grown in the California site in the 2015 season, despite the models not being trained on this data. This results indicate that groups of microbes proliferate predictably between field sites and between years. The sets of OTUs included in these models both increase and decrease in relative abundance over the course of the season (Fig. 3B). The increasing and decreasing OTUs were distinct at the phylum and order levels, suggesting that functional capabilities encoded by these microbes change throughout the season.

These results show that OTUs conserved between two diverse field sites can be used infer the age of rice plants. It is unknown whether our models developed for the rhizosphere and endosphere compartments are generalizable to rice plants grown in other regions of the world: climate, geography, and cultivation practices are all factors which contribute to microbiome structure and would likely effect that age predicting models. The rice rhizosphere microbiota shows similarity at lower resolution taxonomic levels (29), even when grown on different continents, so it is likely that RF models built at the order or family level may allow generalization in predictability across continental scales. Diverse plant species host divergent microbiota assemblages, even at the phylum level (30), thus it is unlikely that our models could be accurately applied to predicting the age of a set of genetically diverse plant species.

To test our hypothesis that environmental variables could affect the accuracy of age prediction, we used the age predicting models to estimate age of plants experiencing drought compared to well-watered conditions (Fig. 4). We found that the predicted age of drought stressed plants using the endosphere models was significantly less than control plants indicating that the water deprived plants were hosting a developmentally immature endosphere microbiota (Fig. 4A). This finding is intriguing given that rice plants experiencing drought stress during the vegetative stage typically have delayed flowering times (31, 32). These data support a model where an arrest in host plant development as a consequence of an environmental disturbance also impacts normal development of the endospheric microbiota. More studies will need to be conducted to confirm this hypothesis. Nonetheless, these results indicate that the models described here can be used to create a baseline for normal microbiota development and to test how environmental or biological perturbations may affect the maturation process. It is unknown how the OTUs included in each model relate to the overall health of the rice plants and assembly of a healthy microbiota. In malnourished Bangladeshi children, the gut microbiota was shown to be underdeveloped compared to healthy children (13). It was found that age discriminant OTUs identified through random forests models from the healthy human gut microbiota prevented growth defects in mice harboring microbiota from malnourished children (33). A similar approach could be used with axenically grown rice plants to identify whether isolated age-discriminant OTUs aid in biotic and abiotic stress relief.

### Late colonizing OTUs show greater conservation between field sites

Over the course of the season, we showed that the rhizosphere and endosphere communities between field sites grow more similar, stabilizing around eight to nine weeks after germination (Fig. 5A). Similarly, there were more age-discriminant OTUs in the rhizosphere and endosphere RF models that showed increasing trends in relative abundance over the course of the growing season (Fig. 3B). These results suggested that there was less conservation between the field sites for early colonizing OTUs. Our data suggests that site-specific OTUs were significantly greater at the beginning of the season in both the rhizosphere and endosphere, but were diminished at later time points in the growing season (Fig. 5B). This effect was greater in the endosphere than the rhizosphere, presumably because the rhizosphere is host to both microbes affected by plant processes as well as microbes from the soil biota that are unaffected by signals originating from the host. The increased conservation of the late-emerging microbiota may be due to plant selectivity, while the early colonizing microbiota may be due to opportunistic colonization of plant tissue by the soil microbiota. Nevertheless, through our random forests approach, we were able to identify specific early colonizing microbes that are shared between the two field sites and these OTUs were almost unanimously enriched in their respective rhizocompartments compared to bulk soil (Sup. Fig. 3), suggesting active or passive recruitment of the conserved early colonizing microbiota. In humans, the early colonizing gut microbiome has been implicated in educating the immune system (34, 35). In plants, it is unclear whether the conserved early colonizing microbiota plays a role in conditioning the activity of the plant innate immune system. A recent study in maize using a synthetic community found that certain bacteria, when omitted from the community, drastically disrupt normal microbiota assembly in roots and that disruption of the normal microbiota assembly led to greater susceptibility to fungal pathogens (36). It may be possible that a portion of the conserved early colonizing taxa may be acting as such keystone species, thus it is of interest to characterize isolated members from the early colonizing age-discriminant OTUs to understand whether they have a role in immune system function and microbiota assembly.

### Advancement of the root-associated microbiota correlates with rice developmental progression

By growing rice varieties with differing developmental trajectories, we were able to quantify the effect of plant development on the root associated microbiota. We observed that there was a gradient in microbiota maturation across the second principal coordinate in the rhizosphere and endosphere, but this effect was absent in the rhizoplane (Fig. 6C). We observed a small but significant differentiation between the tested varieties when using the age-predicting RF models (Sup. Fig. 6), indicating that the OTUs used by these models are primarily effected by plant age rather than developmental stage. By using a Random Forests regression, we were able to identity sets of microbes in each rhizocompartment that are able to distinguish the samples by developmental stage (Fig. 6D). These results suggest that the root microbiota is affected by both plant age and developmental stage and these effects influence different sets of microbes.

Dombrowski et al find no significant differences in microbiota structure between an early flowering *A. alpina* mutant (*pep1*) (37) and non-flowering wild type plant at the same age, suggesting that residence time, rather than developmental stage, is the main driver of microbiota dynamics (15). Our results are not consistent with the Dombrowski et al. study where the microbiome was not affected by plant developmental stage. There are several likely causes for this discrepancy. *Arabis alpina* is a wild perennial plant and rice is a domesticated annual species. One hallmark trait of cereal domestication is the selection for varieties with larger sink sizes in the seed (38). For instance, in wild species, source carbohydrates are more evenly distributed to various sink tissues in the plant than domesticated cereals, where the seeds are a predominant sink (38, 39). The discrepancy between sink-source dynamics in these two host plants could at least partially explain why our study detected shifts in the microbiota due to developmental stage while it was undetected in *A. alpina*. In rice, accumulation and storage of carbohydrates in the stems occur until the onset of reproduction, whereafter internal signals reprogram the host plant to repartition carbohydrates to the developing panicle and to the filling grain (40). These host signals, along with the shifting nutritional needs at this stage could explain the stabilization in the microbiota that occurs at the onset of reproduction. Here we have a provided one of the first links between host plant development and composition of the root-associated microbiota.

### A model for acquisition and dynamics of the root-associated microbiota

Previous studies have implied that the root-associated microbiota is dynamic across the life cycle of the host plant (23–25, 41), but due to low temporal resolution of sampling, the patterns of compositional changes have not been elucidated. Together with these studies, our findings help to paint a clearer picture of the dynamics of the root-associated microbiota over the lifecycle of rice plants. We propose a two-step model to explain this process over the life cycle of annual plants: Immediately after germination, a community of early colonizing microbes establish in and around the roots. It is unclear to what extent the host actively drives this process as many of the early acquired microbes are site-specific and may represent opportunistic colonization by a subset of the soil microbiota. Nevertheless, a set of early colonizing microbes conserved between field sites were identified using our RF models and may represent consortia that are responsive to cues from the host plant. At around the time of entering the reproductive phase, a later colonizing set of microbes displaces the early colonizing microbes which then remains stable throughout the remainder of the life cycle of the host plant. The rate of these transitions is dependent upon life cycle progression trajectories of different genotypes. The microbes colonizing later in development were more conserved between the field sites, which are perhaps influenced by factors from the host. Root exudate composition varies significantly over the life cycle of Arabidopsis and tomato (42, 43), offering one potential explanation for root microbiota dynamics. Similarly, cereal crops display decreased susceptibility to some pathogens during the adult phase (44–46), suggesting that immune system activity is dynamic across the life cycle of the host plant and may in turn modulate dynamics of the root microbiota, explaining the turnover of microbes between the vegetative and reproductive stage. The plant immune system is likely to play a larger role in affecting dynamics of the rhizoplane and endosphere microbiota than the rhizosphere as microbes on the root surface and interior make physical contact with the host plant cells. This hypothesis is supported by our findings, where we see a more pronounced stabilization effect in the rhizoplane and endosphere than the rhizosphere (Fig. 1D).

Because this study is the first high-resolution study examining the dynamics of the plant microbiota, it remains unknown whether similar stabilization dynamics exist in microbiota acquisition across other host plant species. It may be observed that stabilization in other plant species takes a shorter or longer period of time depending on the length of life cycle of the host. Similarly, it is of interest to study root microbiota dynamics in perennial plants over the course of several years to observe whether the microbiota reaches steady state at maturity or whether the microbiota requires re-establishment in each subsequent growth season. In addition, it is of interest to understand the functional and transcriptional activity of the dynamic microbiota to unravel plant-microbe communication throughout this process.

Manipulation of the soil microbiota using bacterial isolates to increase crop yield and resistance to pathogens has been proposed to be an important feature of the next Green Revolution (47). Using mono-association assays, arrays of plant growth promoting or disease suppressing bacteria have been identified (48, 49). However, a major problem facing seed companies, agronomists, and researchers is inconsistent persistence of plant beneficial bacteria once inoculated into complex soil microbial communities (50–52). Our findings indicate that root-associated microbiota are sensitive to plant age and development. These results imply that inoculation of plant growth promoting bacteria should be age-appropriate for the host. For instance, a plant growth promoting bacterium which normally establishes during the vegetative phase would not be a good candidate for coating onto seeds because it may not survive during the seedling stage. Using the experimental and analytical methods outlined in our study will allow researchers and seed companies to better understand when and how potential plant growth promoting bacteria should be applied to the crop species of choice.

## Materials and Methods

### Plant growth and sampling during the California 2014 and 2015 seasons

We collected samples from a commercially cultivated rice field in Arbuckle, CA. In both seasons the field was water seeded in early May. Water seeding is conducted by first preparing the field by removing winter vegetation, disking the soil to produce baseball sized clods, applying nutrients, and then flooding. The rice seeds are soaked in water overnight and then loaded into an aircraft where they are then applied aerially to the field in an even density. For this particular field, the farmer grew the M206 cultivar, a medium grained California variety that has an average heading date of 86 days after germination. We began sampling plants 7 days after the fields were seeded, which coincided with the emergence of the seminal roots. In the 2015 and 2016 seasons we restricted our area of sampling to a 150 x 150 foot section of the field. Within this section we sampled plants from random locations. Our sampling occurred as previously described (6). Using gloved hands, we would scoop under the root mass to separate the plant from the ground. Grabbing the plant by the base of the stem, we would then shake the plant to remove loosely attached soil from the roots. We would then place the roots with tightly adhering soil into 50 mL Falcon tubes with 15 mL autoclaved phosphate buffered saline (PBS) solution. We would then bring the samples back to the laboratory at UC Davis for subsequent processing.

### Plant growth during the Arkansas 2016 Season

In the Arkansas 2016 season, we grew the tropical japonica variety, Sabine, and a hybrid variety, CLXL745, in a split-plot design in an agricultural field near Jonesboro, AR. Each plot was isolated from other plots via berms and each plot had its own source of water and drainage (see design, Sup. Fig. 8). The roots of the plants were collected over the growing season as described above, placed into 50 mL Falcon tubes, and shipped overnight on ice to UC Davis. At UC Davis, the root-associated compartments were separated (see below) and stored at -80°C until DNA could be extracted.

### Plant growth during the California 2016 Season

We grew closely related temperate japonica cultivars in the same Arbuckle, CA field for the 2016 season. Kitaake is a variety typically used under laboratory settings due to its relatively fast life cyle. M206 and M401 are varieties adapted to growing in California with medium times to panicle initiation. M401, however, requires a longer period of time for flowering. Nipponbare is a variety with a longer time to panicle initiation and flowering. Briefly we designed a fully randomized block design to grow these varieties in 1 x 1 m plots in quadruplicate (see design Sup. Fig. 9). We left 0.5 m wide walking lanes between each plot which subsequently allowed us to sample bulk soil throughout the course of the growing season. To be consistent with the previous years’ methods, we water seeded these varieties. This required soaking the seeds in a 2% bleach solution for 4 hours to remove the risk of the field being contaminated *Fusarium moniliforme*, a seed borne fungal pathogen that causes the disease Bakanae (http://ipm.ucanr.edu/PMG/r682100711.html). The seeds were then washed 3 times with sterile water and soaked overnight. We then hand seeded each plot at similar density as what the farmer had applied in previous seasons. At each time point the plants were sampled as previously described and transported to the lab for further processing. The plants during this season were dissected to ascribe various developmental stages in accordance with Counce et al. (53). Because the rice plants were water seeded, there was a high chance that each genotype’s plot could be contaminated by seeds drifting in from another plot or seeds that the field manager had planted. The genotypes were confirmed by amplifying SSR marker RM144 (54) using the endosphere DNA samples collected from the plants. From this analysis, we excluded the first week of Nipponbare samples from the analysis due to all of the samples being contaminated by M206. Similarly, we removed other samples from the analysis where the SSR marker genotyping did not match the genotype of the plot in the field.

### Separation of the root-associated compartments and DNA extraction

In each instance of sampling roots, we collected material from the first inch of roots just below the root-shoot junction. _The root associated compartments were separated as previously described (6). The roots with soil attached were vortexed in PBS solution and 500 uL of the resulting slurry was used for DNA extraction. The roots were cyclically washed in fresh PBS solution until no soil particles were visible in the solution. The roots were then placed into fresh PBS and sonicated for 30 s to remove surface cell layer of the roots. The resulting slurry was centrifuged down to concentrate the biomass and then used as the rhizoplane fraction for DNA extraction. The remaining roots were sonicated twice more, refreshing the PBS solution at each stage, and then ground in a bead beater. The resulting solution was used for DNA extraction as the endosphere fraction. All DNA extraction was performed using the MoBio Powersoil DNA isolation kits.

### PCR amplification and sequencing

We amplified the V4 region of the 16S ribosomal RNA gene using the universal 515F and 806R PCR primers. Both our forward and reverse primers contained 12 base pair barcodes, thus allowing us to multiplex our sequencing libraries at over 150 libraries per sequencing run (6). Each library was accompanied by a negative PCR control to ensure that the reagents were free of contaminant DNA. We purified the PCR products using AMPure beads to remove unused PCR reagents and resulting primer dimers. After purification, we quantified the concentration of our libraries using a Qubit machine. Our libraries were then pooled into equal concentrations into a single library and concentrated using AMPure beads. The pooled library then went through a final gel purification to remove any remaining unwanted PCR products. Pooled libraries were sequenced using the Illumina MiSeq machine with 250 x 250 paired end chemistry.

### Rice Drought Microbiome Data

Data for the rice drought microbiome experiment performed by Santos-Medellín et al. was recovered from the NCBI Short Read Archive using project number <u>PRJNA386367</u>. All sequences were filtered and clustered in accordance with the other MiSeq derived sequences (see below).

### Sequence Processing, OTU clustering, and OTU filtering

The resulting sequences were demultiplexed using the barcode sequences by in house Python scripts. The sequences were quality filtered and then assembled into full contigs using the PandaSeq software (55). Any sequences containing ambiguous bases or having a length of over 275 were discarded from the analysis. The high-quality sequences were clustered into operational taxonomic units (OTUs) using the Ninja-OPS pipeline (56) and then assembled into an OTU table. This OTU table was filtered to remove plastidial and mitochondrial OTUs and filtered to remove OTUs that occur in less than 5% of the samples. This process removed the total OTU count from 24,048 to 8,554 taxa. The resulting 8,554 taxa were used for the analysis.

### Statistical Analysis

All statistical analyses were conducted using R version 3.1 (57). Unless otherwise noted, we determined statistical significance at α = 0.05 and, where appropriate, corrected for multiple hypothesis testing using the Bonferroni method. Shannon diversity was calculated using the diversity() function, PCoA analyses were conducted using the capscale() function, PERMANOVA was conducted using the adonis() function from the Vegan package (58). Linear models were run using the lm() function and ANOVA was run using the aov() function. Beta regression was performed using the BetaReg package (59). Random forests models were generated and analyzed using the RandomForests package (60). All graphs and plots were generated using the ggplot2 package (61). Differential OTU abundance was performed using exact tests in the package edgeR (19). R notebooks for the full analyses can be found at https://github.com/bulksoil/LifeCylceAnalysis.

### Accession Numbers

All data will be submitted to the NCBI short read archive at a future date.

---

**Supplemental Figure 1.**
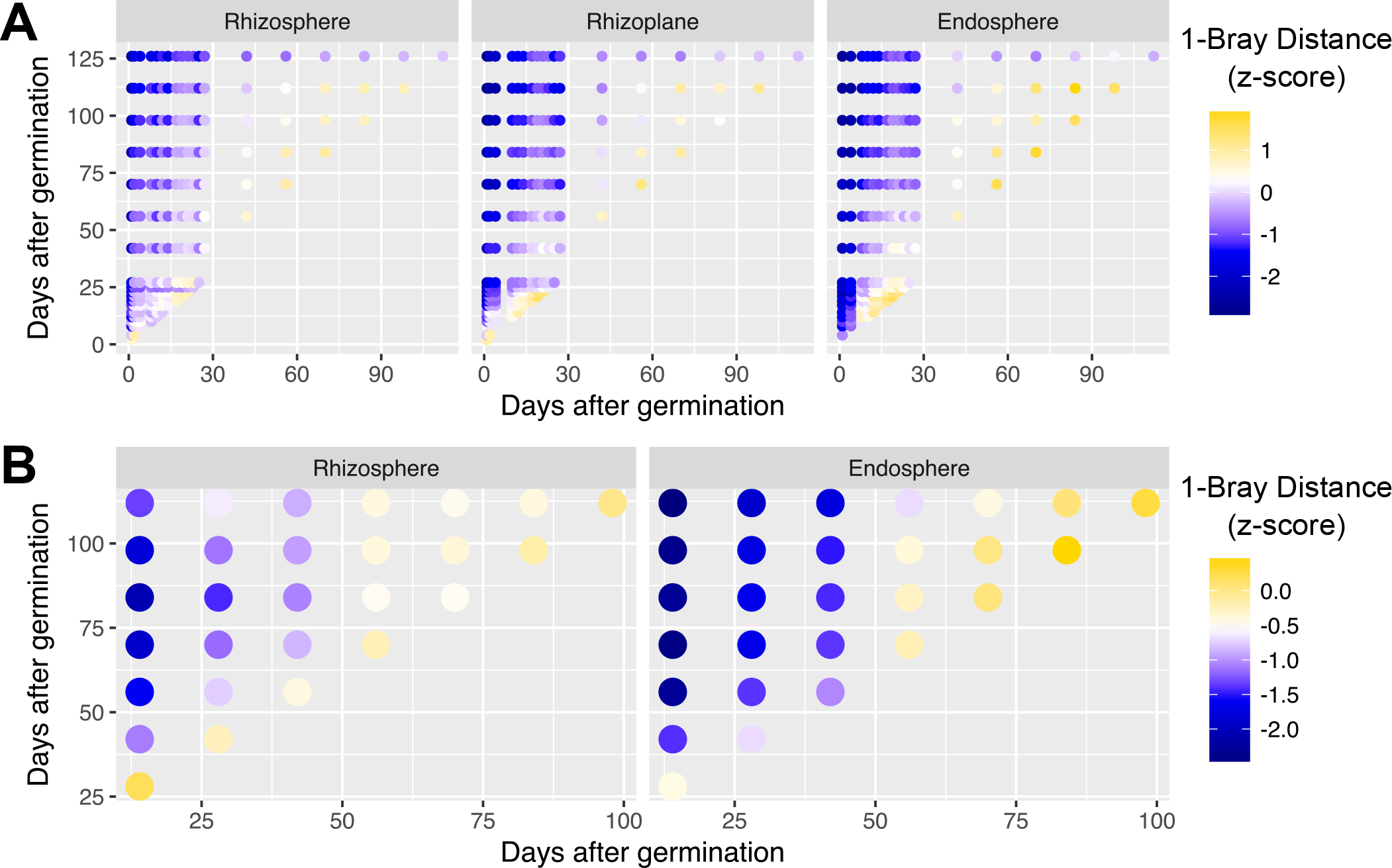
Stabilization in microbiome dynamics occurs in multiple sites and seasons. **A)** Heatmap of pairwise similarities between time points for the California field site in the 2015 season. **B)** Heatmap of pairwise similarities between time points for the Arkansas field site in 2016.

**Supplemental Figure 2.**
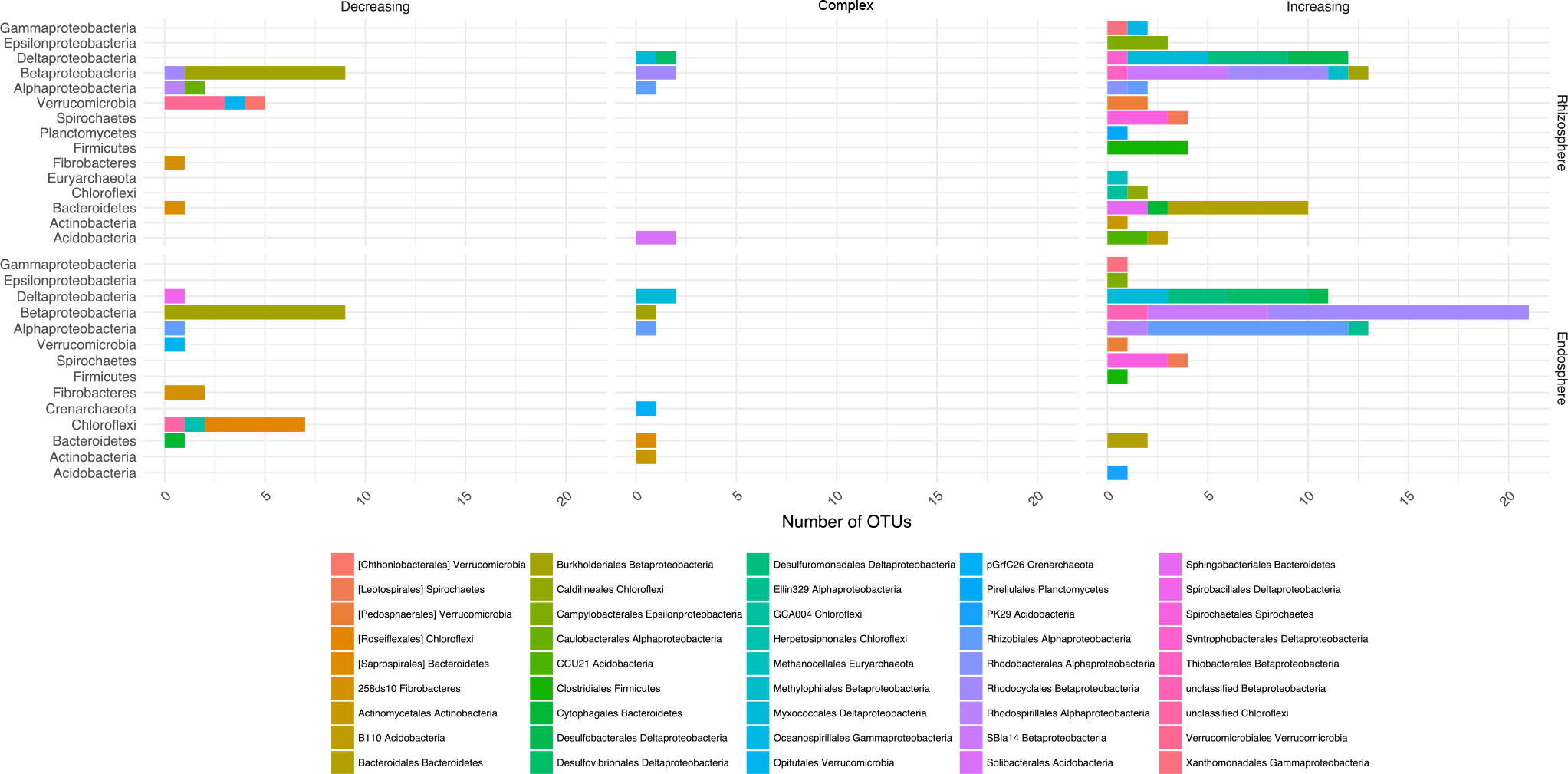
Decreasing, complex, and increasing age discriminant OTUs are divergent at the order level.

**Supplemental Figure 3.**
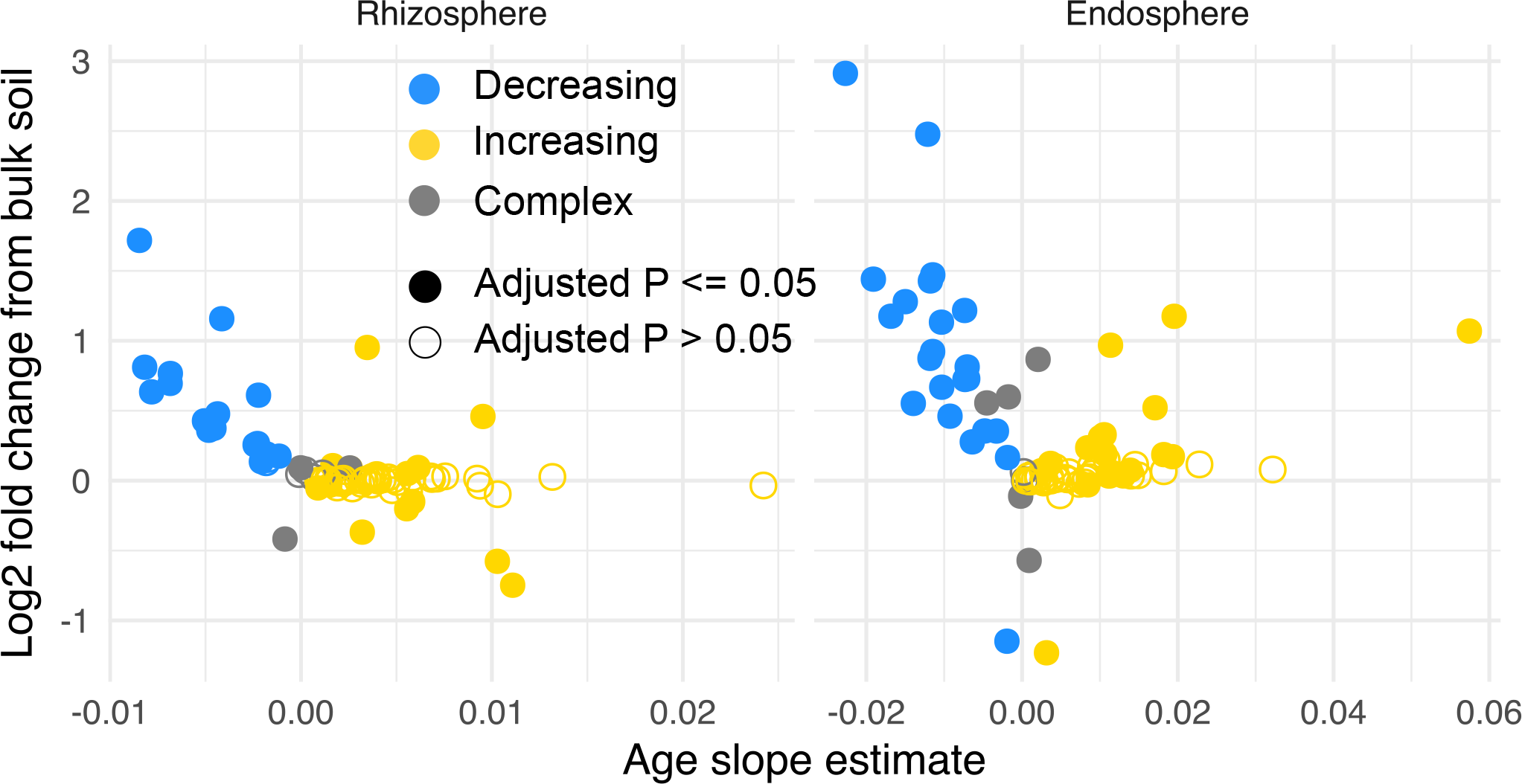
Age discriminant OTUs with decreasing abundance over time are enriched in their respective rhizocompartments compared to bulk soil.

**Supplemental Figure 4.**
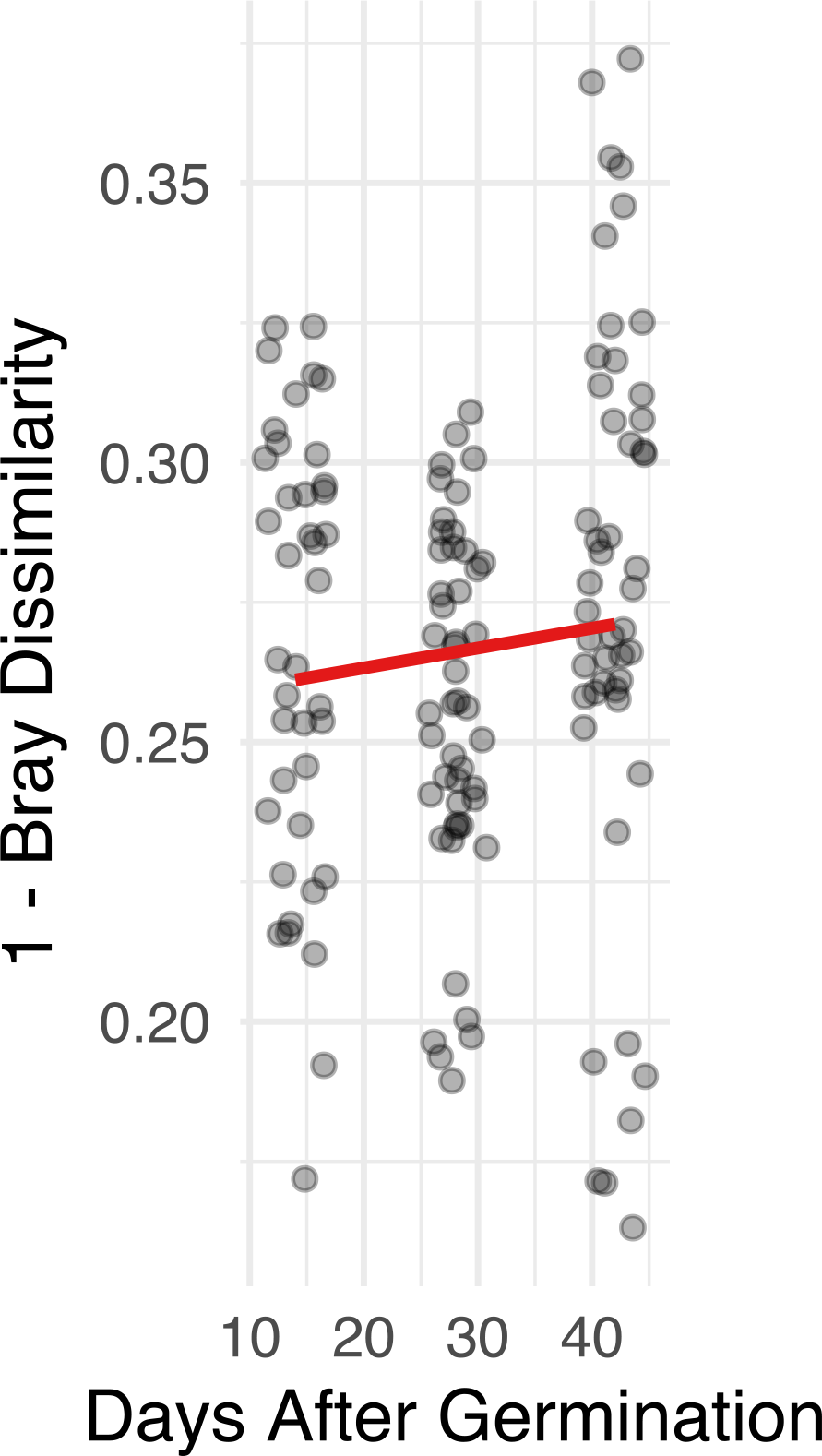
Bulk soil communities do not become more similar between field sites over time. Pairwise Bray dissimilarity measures between bulk soil samples within the same age across the Arkansas 2016 and California 2014 field sites.

**Supplemental Figure 5.**
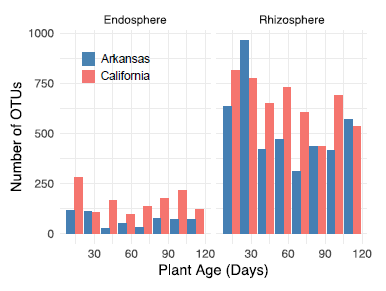
California 2016 samples predicted by the sparse age-discriminant RF models. **A)** Age predictions for the rhizosphere and endosphere using the sparse RF models. **B)** Relative abundance of increasing/complex/decreasing age-discriminant OTUs.

**Supplemental Figure 5.**
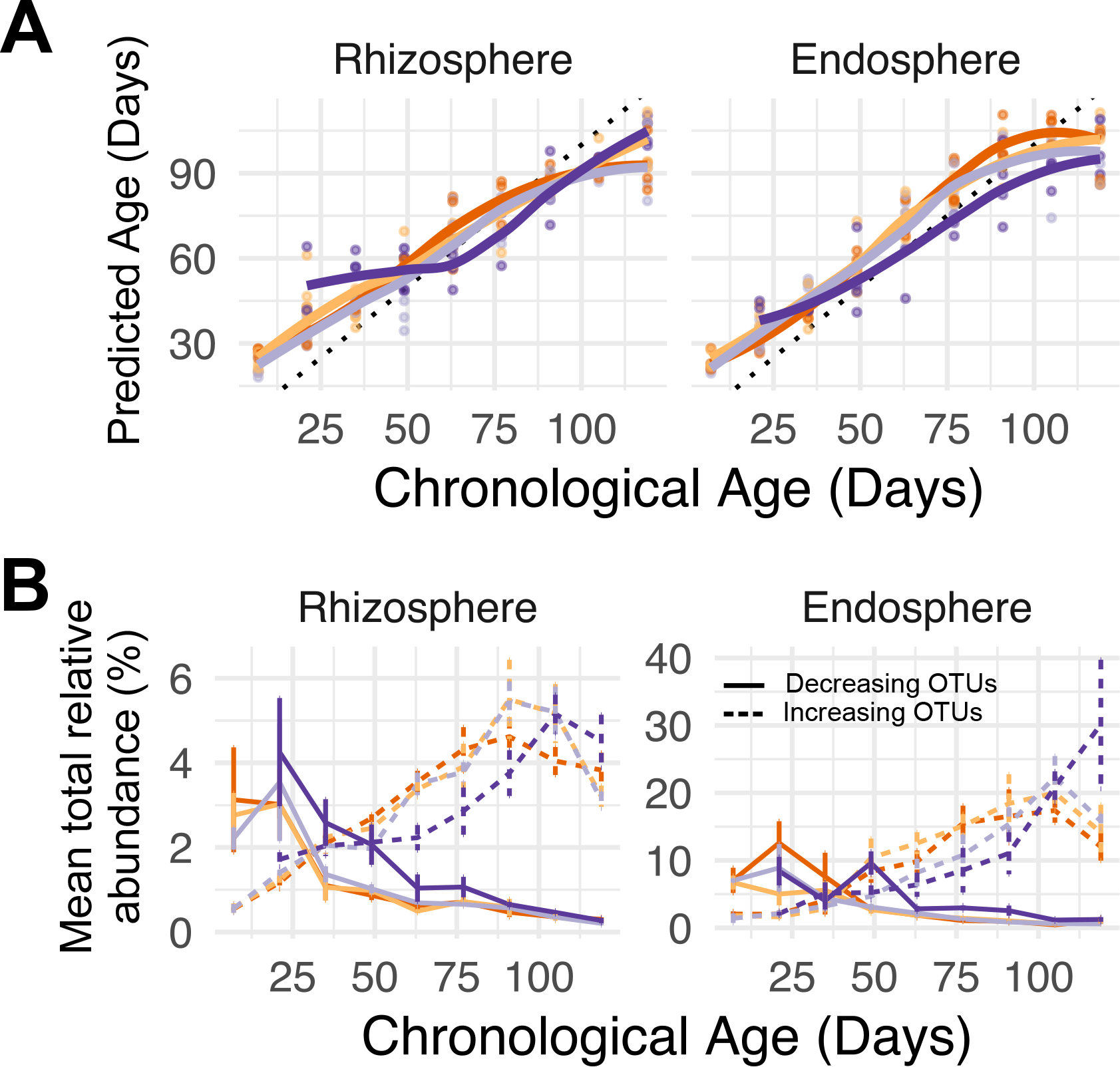
Phylum and Order distribution of the developmental stage discriminant random forests models.

**Supplemental Figure 6.**
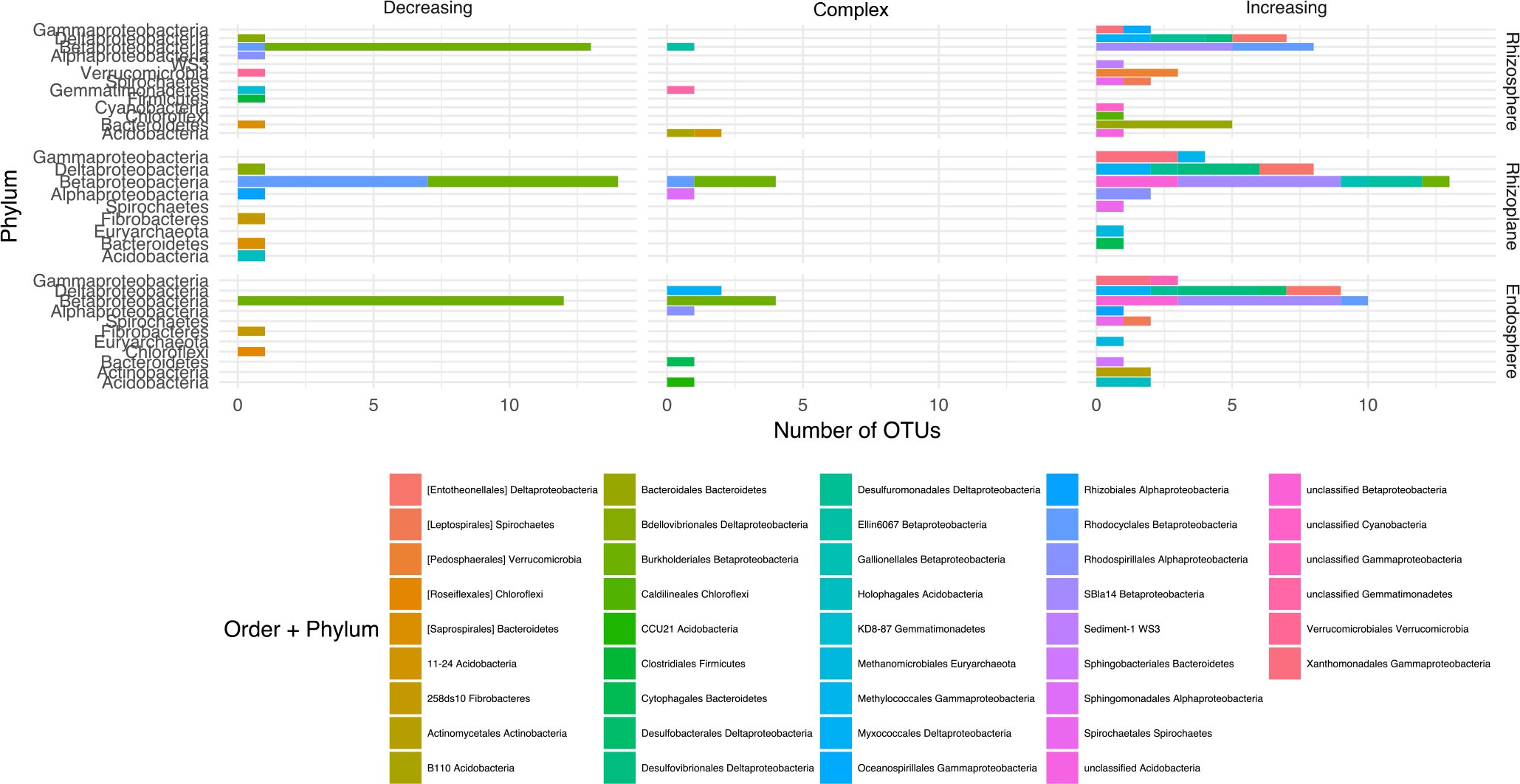
Layout for the Arkansas 2016 field study.

**Supplemental Figure 7.**
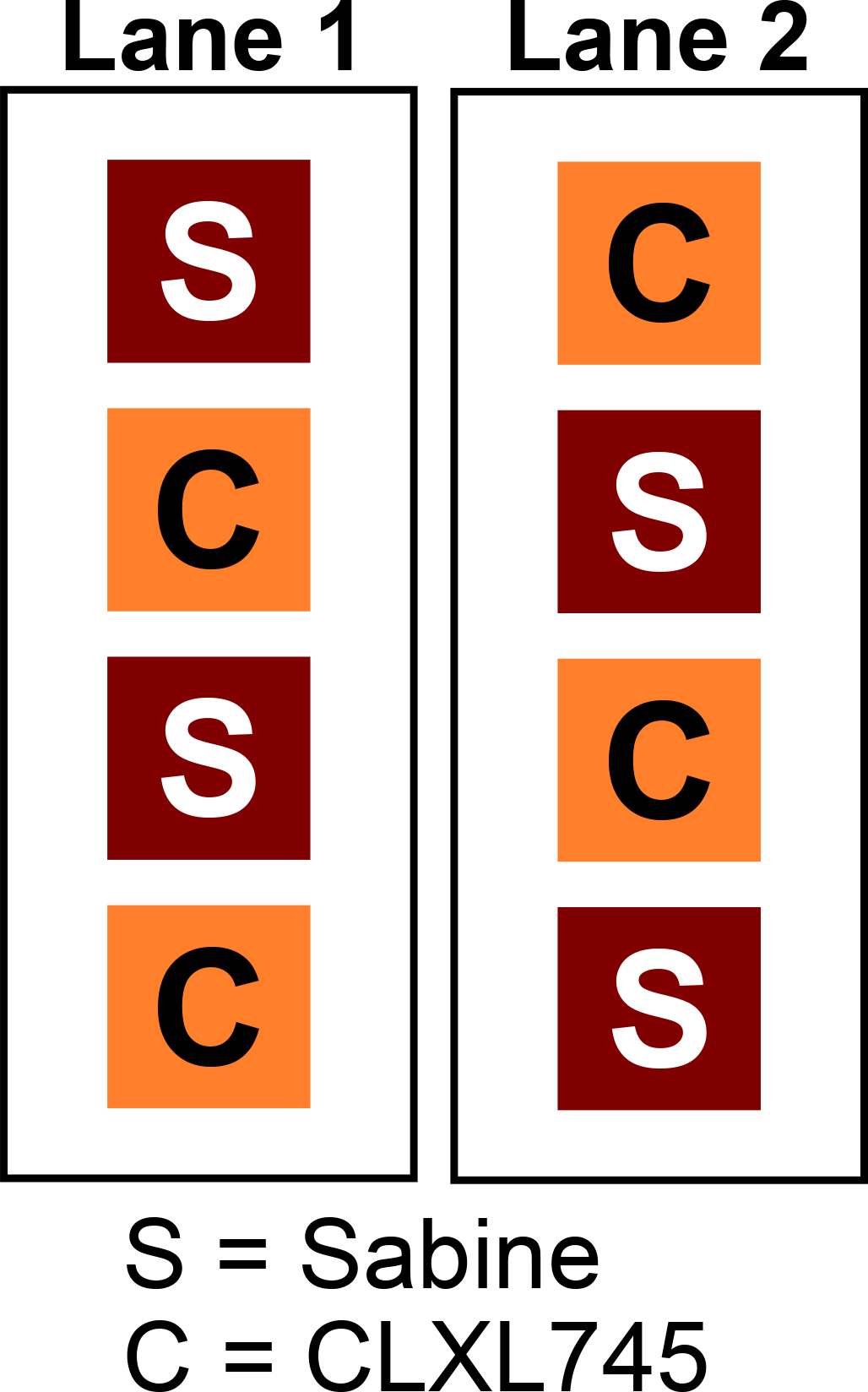
Layout for the California 2016 field study.

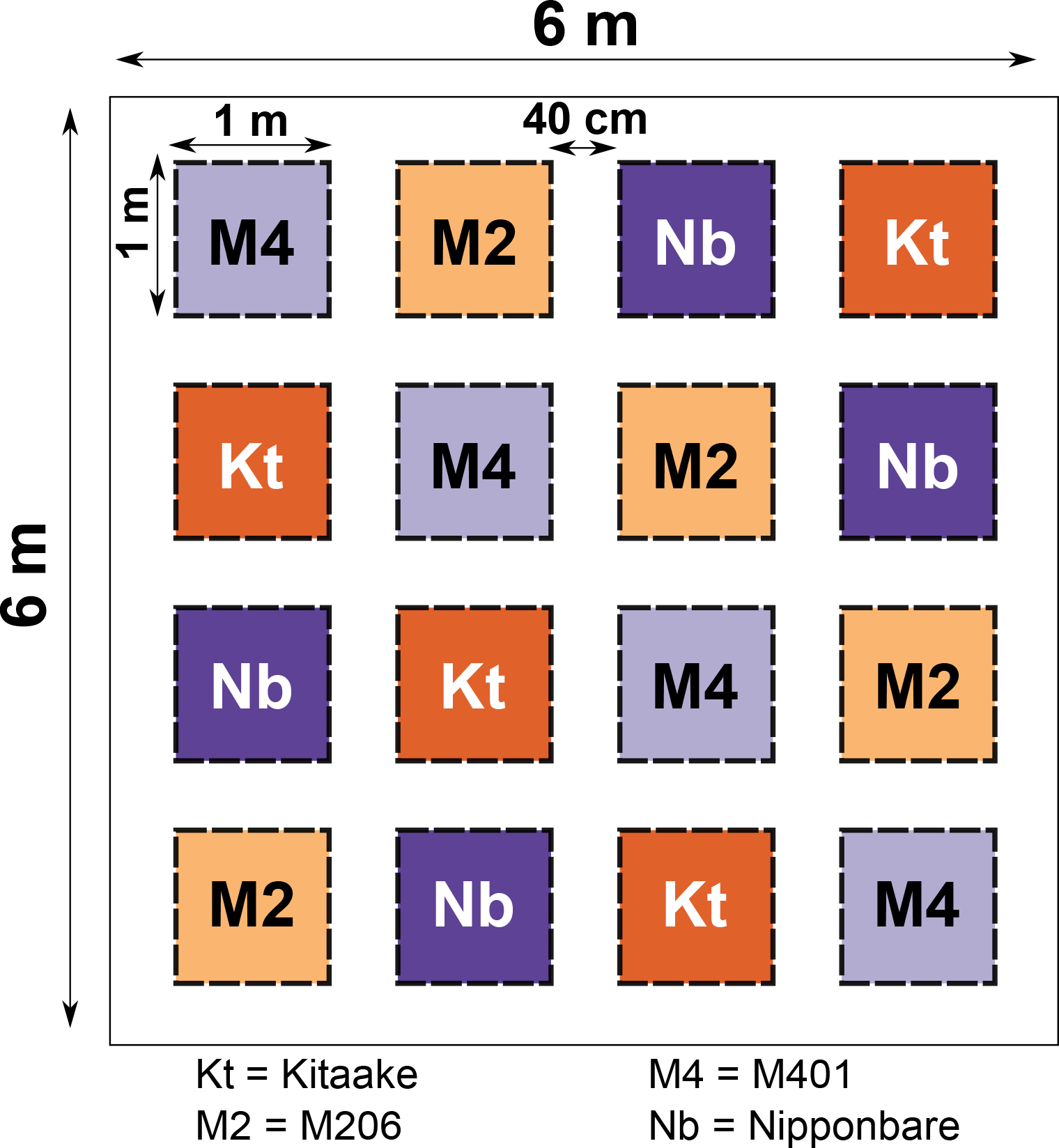

